## Supplemental Figures (Figure S1 to Figure S17) for "Single-cell analysis of mosquito hemocytes identifies signatures of immune cell subtypes and cell differentiation" (from another version of this preprint: version 1, 2020-07-11)

### Supplemental Information

### Supplemental Figure Legends

**Figure S1.** Quality assessment of scRNA-seq analysis on mosquito immune cells.

**Figure S2.** FACS sorting of mosquito immune cell populations.

**Figure S3.** Naïve and blood-fed immune cells isolated in FACS analysis.

**Figure S4**. Comparisons of immune cell clusters to the *An. gambiae* phagocyte proteome.

**Figure S5.** Comparisons of immune cell clusters to previously defined PPO6$^{low}$ and PPO6$^{high}$ hemocyte populations.

**Figure S6.** Expression of mosquito immune genes across cell clusters.

**Figure S7.** Expression of serine protease inhibitors (SRPNs) and CLIP serine proteases across cell clusters.

**Figure S8.** Expression of chemosensory genes across cell clusters.

**Figure S9.** Expression of specific tRNAs across cell clusters.

**Figure S10.** RNA-FISH of *SCRB9*$^{+}$ immune cells.

**Figure S11.** Expression profile and RNA-FISH of *SCRB3*$^+$ immune cells.

**Figure S12.** Co-hybridization of SCRB9 and LRIM15 labels distinct immune cell populations by RNA-FISH.

**Figure S13.** Expression of granulocyte and oenocytoid markers to distinguish immune cell sub-types.

**Figure S14.** Marker gene expression across immune cell clusters used to define Monocle3 lineage analysis.

**Figure S15.** Enriched transcripts associated with the lineage progressions of mosquito immune cells.

**Figure S16.** Phagocytic properties of a subset of *lozenge (lz)*$^+$ cells.

**Figure S17.** Validation of *lozenge* knockdown following RNAi.

### Supplemental Tables

**Supplemental Table S1.** FKPM values of individual immune cells following scRNA-seq analysis.

**Supplemental Table S2.** Differential gene expression in immune cell clusters displaying significant differences between naive and blood-fed cells.

**Supplemental Table S3.** Genes expressed in more than >80% of cells within each respective immune cell cluster.

**Supplemental Table S4.** Averaged gene expression of cells within each immune cell cluster.

**Supplemental Table S5.** Gene subsets identified by Monocle3 analysis.

**Supplemental Table S6.** Primers for qRT-PCR and dsRNA-mediated gene silencing.

**Supplemental Table S7.** Primers for RNAi.

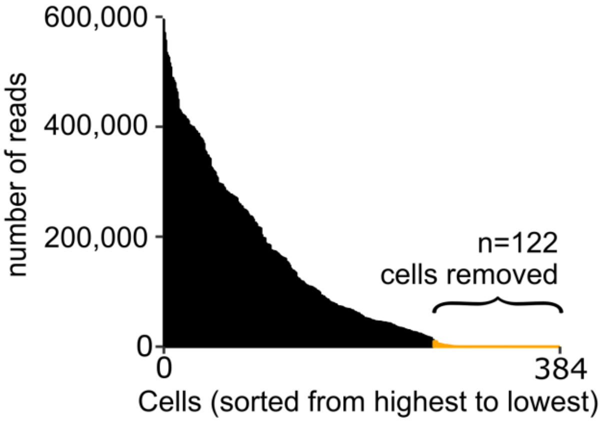


**Figure S1. Quality assessment of scRNA-seq analysis on mosquito immune cells.** The number of transcript reads per cell was used to determine the quality of single-cell data. Using a cutoff of 10,000 reads/cell, we removed 122 cells from our analysis, resulting in a total of 262 cells.

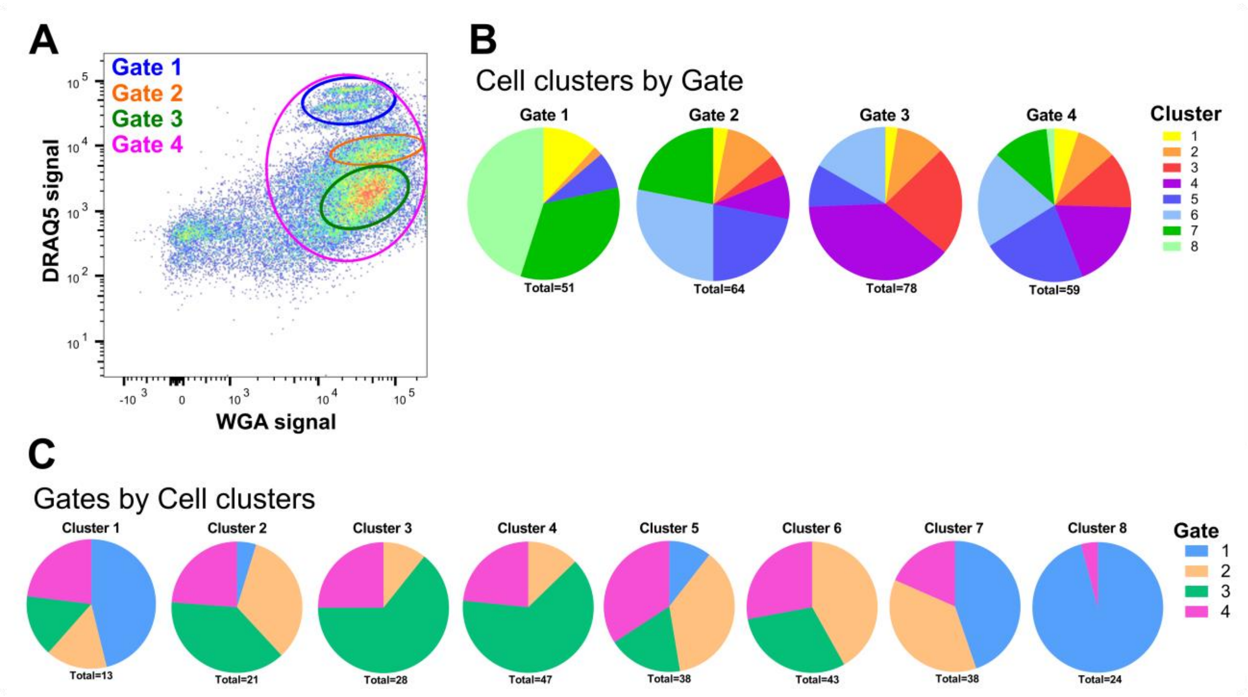


**Figure S2. FACS sorting of mosquito immune cell populations. (A)** Representative image of *An. gambiae* immune cell populations examined by flow cytometry. Distinct sub-populations of cells were separated into four gates: Gates 1-3 as unique populations based on WGA and DRAQ5 signal and Gate 4 to sample a mixed cell population. **(B)** The unique cell clusters identified in our RNA-seq analysis are visualized by the FACS gating from which they originated. **(C)** An additional breakdown of immune cell clusters by gate.

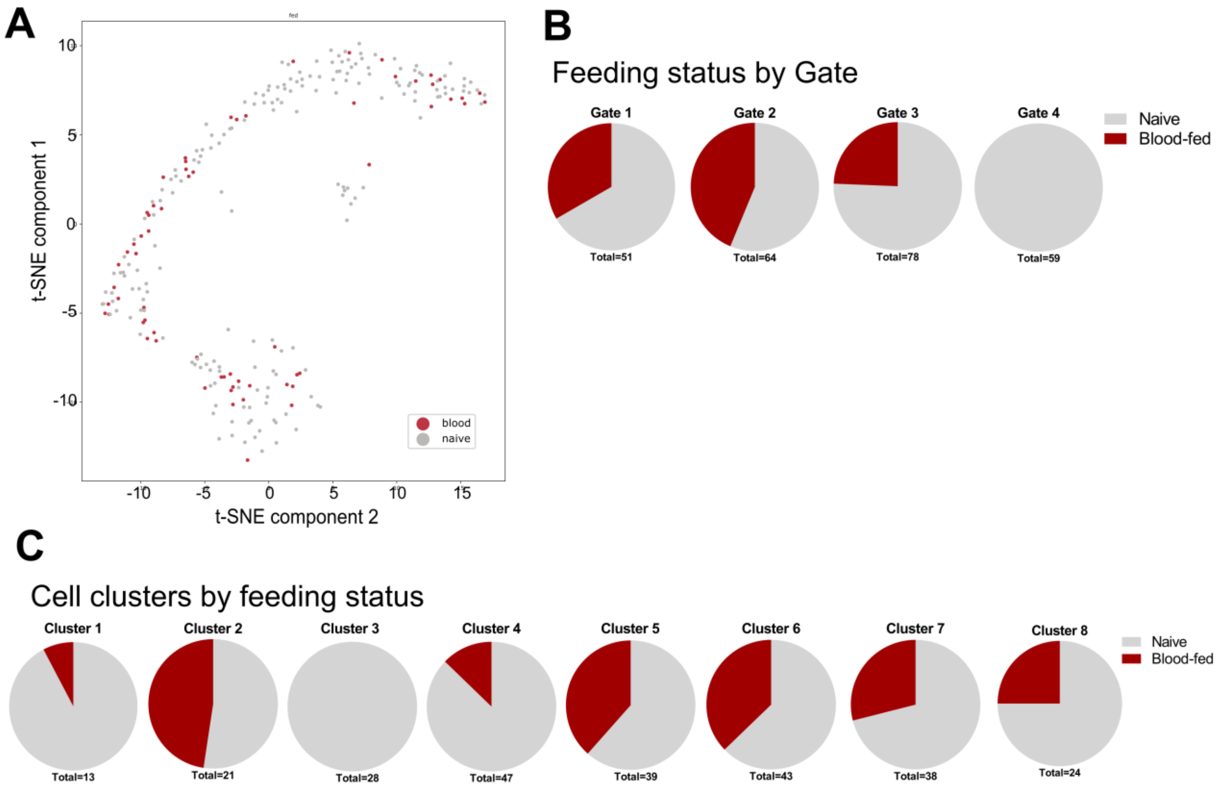


**Figure S3. Naïve and blood-fed immune cells isolated in FACS analysis. (A)** Mosquito immune cells isolated from naïve or blood-fed conditions visualized by t-SNE. **(B)** Cells from naïve and blood-fed conditions were isolated in each of the FACS gatings, except for the presence of blood-fed samples in Gate 4 which were not included in our methodology. **(C)** An additional breakdown of immune cell clusters by feeding status reveals the presence of these immune cell populations under both naïve and blood-fed conditions, with the exception of Cluster 3 that was only identified under naïve conditions.

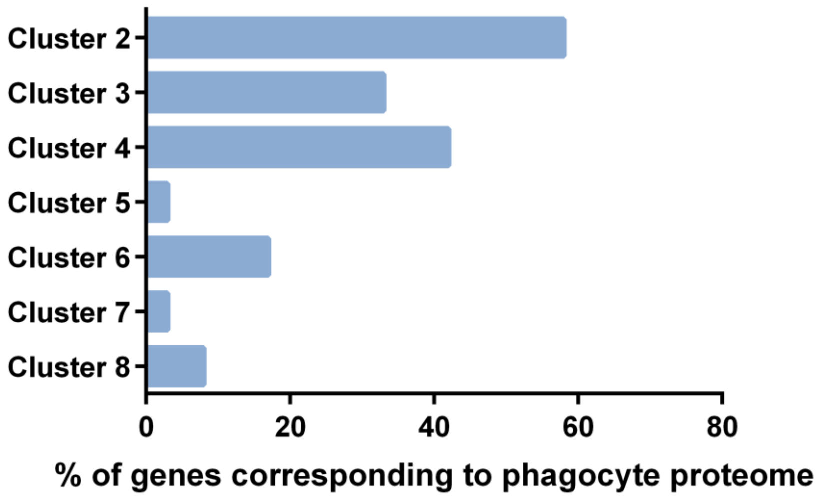


**Figure S4. Comparisons of immune cell clusters to the *An. gambiae* phagocyte proteome.** Transcripts expressed in >60% of cells of each immune cell cluster were compared to the 811 proteins identified in the naïve phagocyte-enriched proteome previously published by Smith *et al.* 2016. Data are displayed as the percentage of genes identified in the proteome with corresponding transcripts from each respective immune cell cluster.

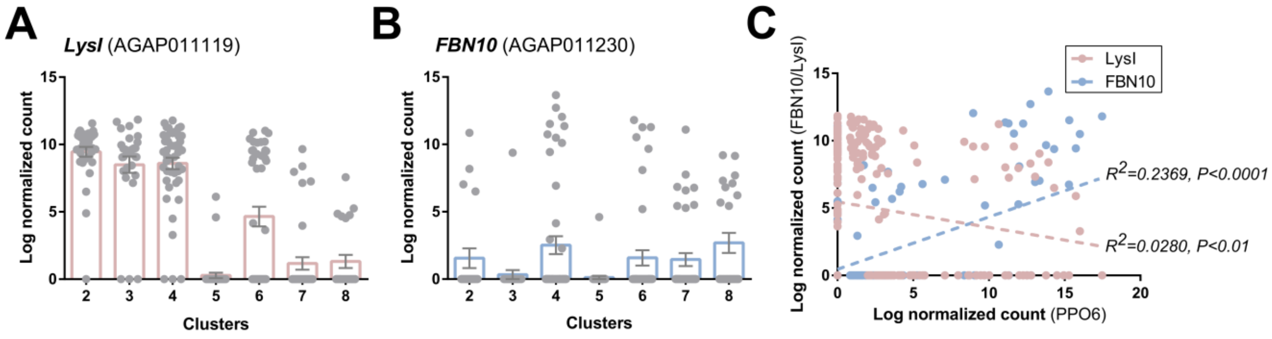


**Figure S5. Comparisons of immune cell clusters to previously defined PPO6$^{low}$ and PPO6$^{high}$ hemocyte populations.** To correlate the immune cell clusters identified in our analysis with previously described PPO6$^{low}$ and PPO6$^{high}$ hemocyte populations (Severo et al. 2018), we examined the expression of *LysI* **(A)** and *FBN10* **(B)** as respective markers of these previously defined cell populations. Although the expression of *LysI* and *FBN10* was distributed across multiple cell clusters in our analysis, transcripts displayed strong negative or positive correlations respectively to the expression of PPO6 **(C)**.

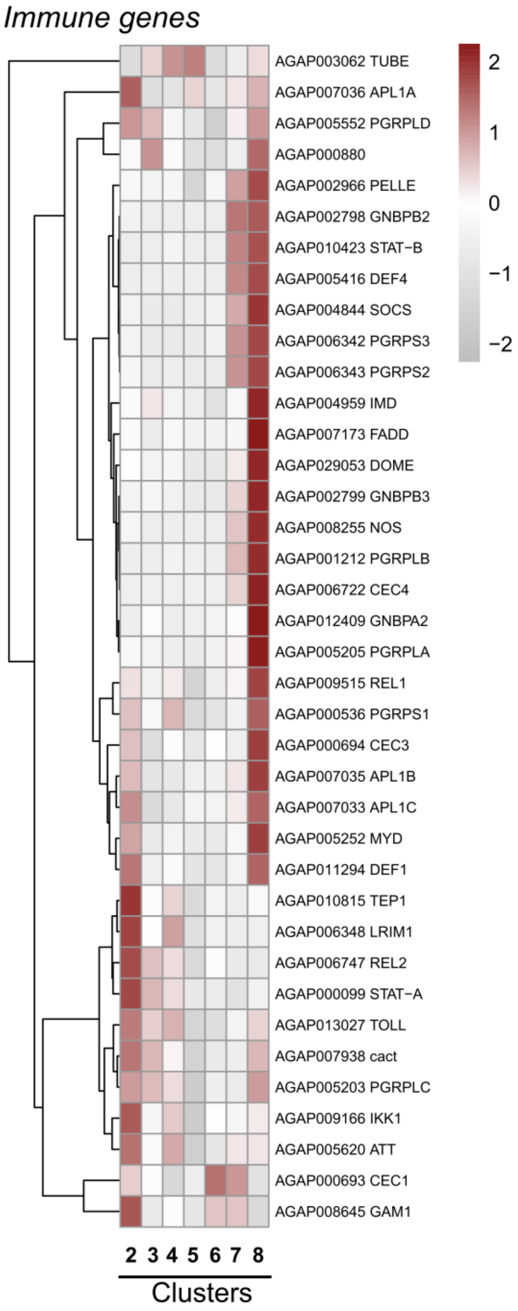


**Figure S6. Expression of mosquito immune genes across cell clusters.** Expression patterns of known components of the *An. gambiae* IMD, Toll, and JAK-STAT pathways, as well as other antimicrobial proteins (AMPs) and complement factors with integral roles in mosquito innate immunity displayed across immune cell clusters.

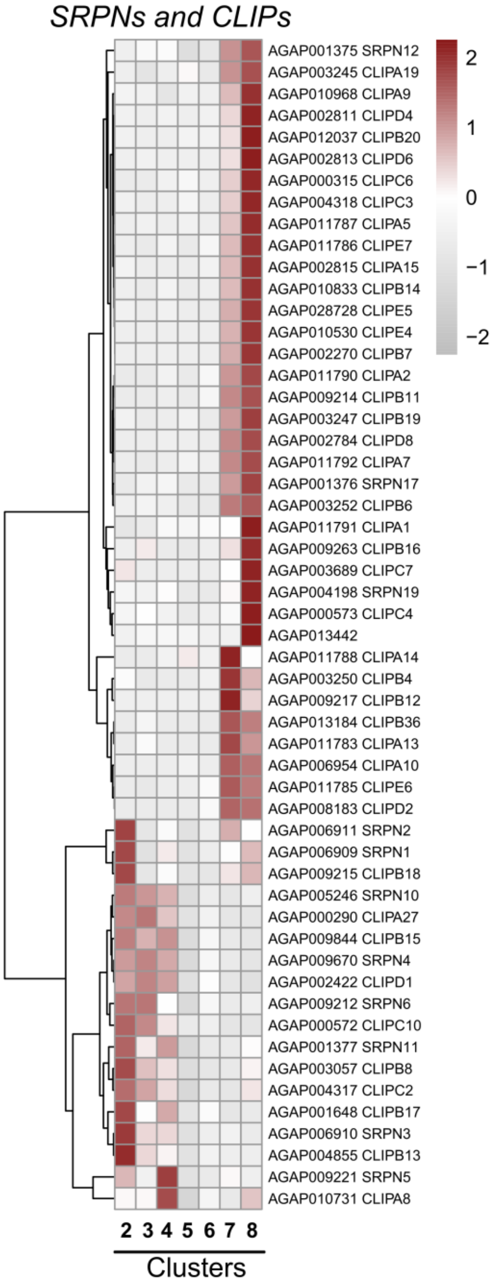


**Figure S7. Expression of serine protease inhibitors (SRPNs) and CLIP serine proteases across cell clusters.** Expression patterns of SRPN and CLIP family members across immune cell clusters.

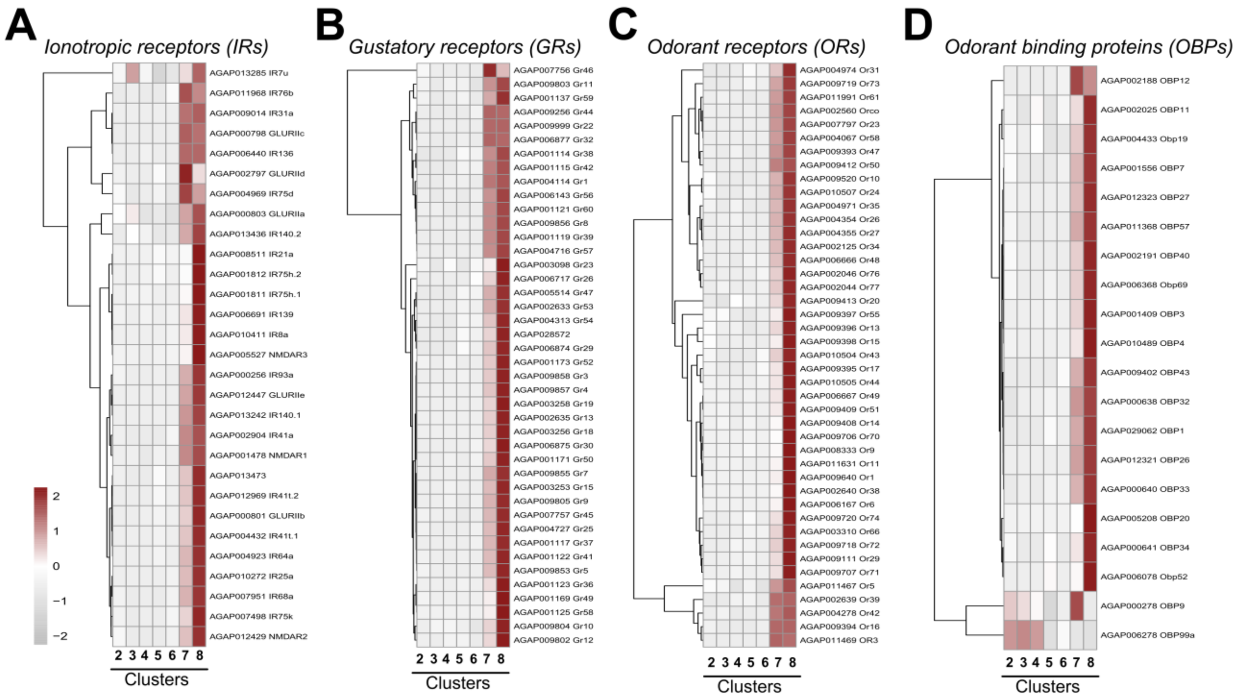


**Figure S8. Expression of chemosensory genes across cell clusters.** Expression patterns of annotated ionotropic receptors (IRs) **(A)**, gustatory receptors (GRs) **(B)**, odorant receptors (ORs) **(C)**, and odorant binding proteins (OBPs) **(D)** across immune cell clusters.

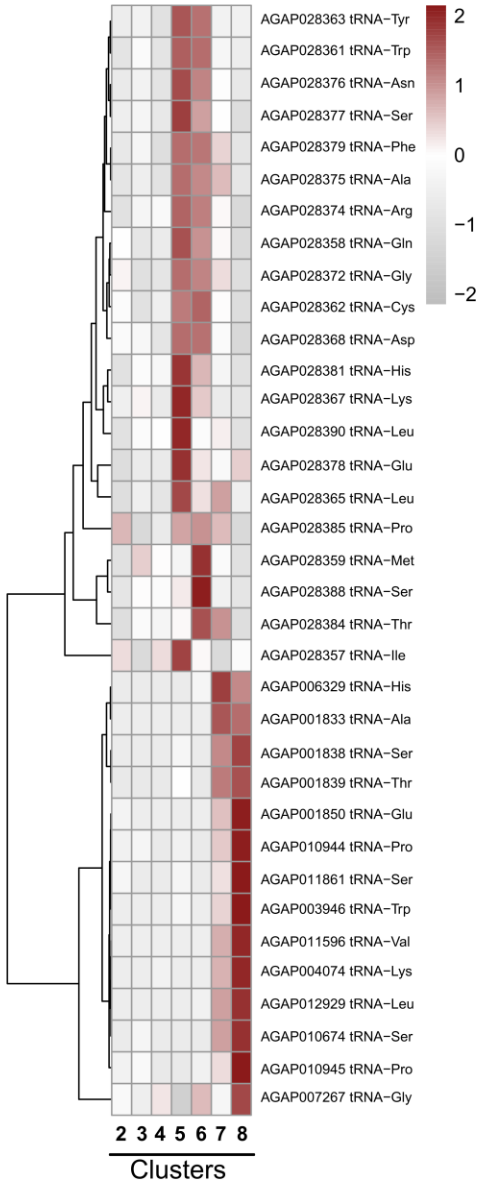


**Figure S9. Expression of specific tRNAs across cell clusters.** Expression patterns of specific tRNAs across immune cell clusters.

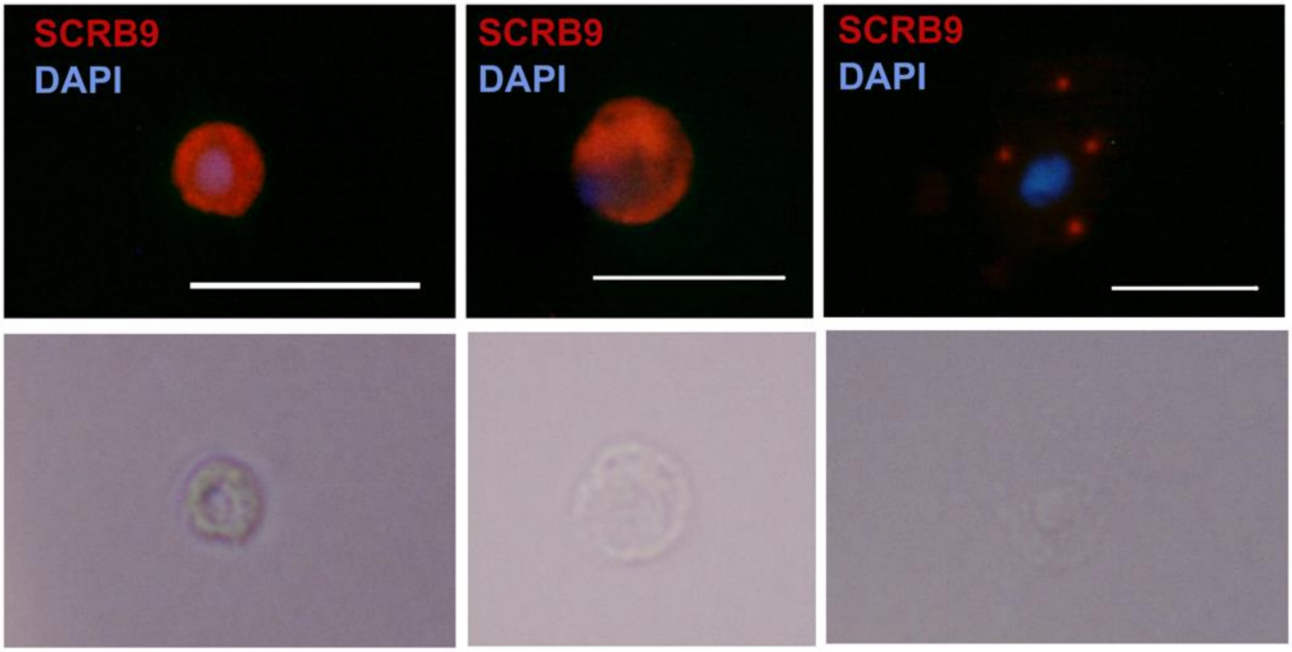


**Figure S10. RNA-FISH of *SCRB9*⁺ immune cells.** RNA-FISH images of immune cells labeled with SCRB9 and stained with DAPI. Below is the corresponding phase contrast image of the same fixed cell. Scale bar, 10 µm.

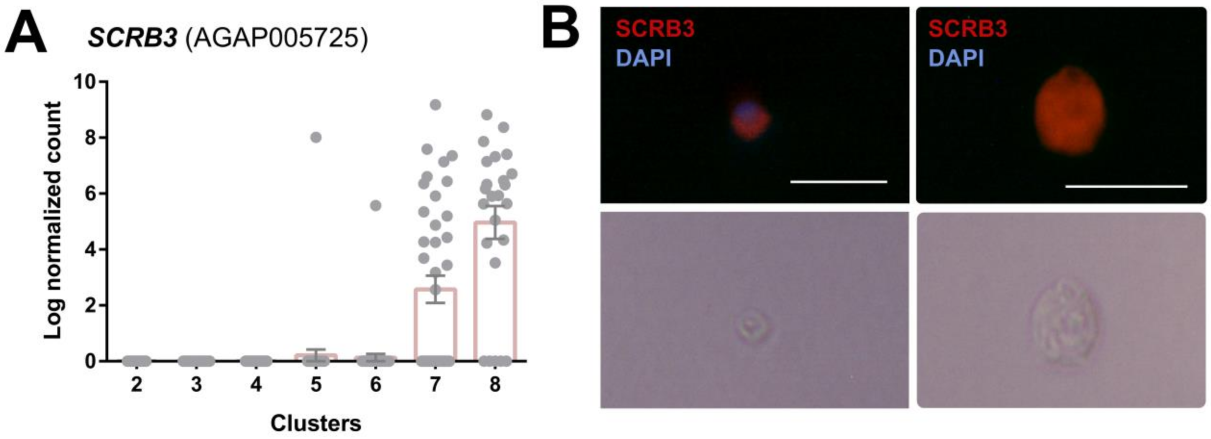


**Figure S11. Expression profile and RNA-FISH of *SCRB3*⁺ immune cells. (A)** Expression of *SCRB3* across immune cell clusters. **(B)** RNA-FISH images of immune cells labeled with SCRB3 and stained with DAPI. Below is the corresponding phase contrast image of the same fixed cell. Scale bar, 10 μm.

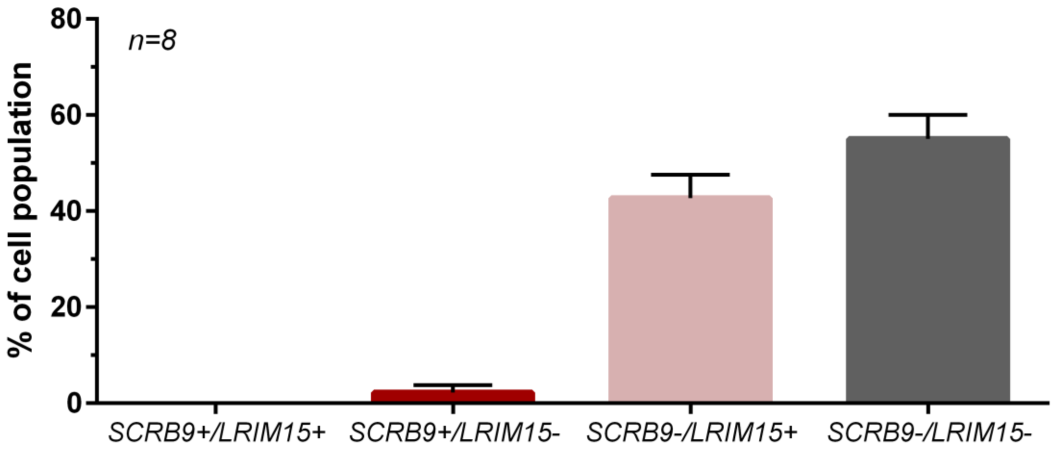


**Figure S12. Co-hybridization of SCRB9 and LRIM15 labels distinct immune cell populations by RNA-FISH.** RNA-FISH experiments were performed with both SCRB9 and LRIM15 probes, with the resulting percentage of cells expressing either, both, or neither immune cell marker. n, number of individual mosquitoes examined.

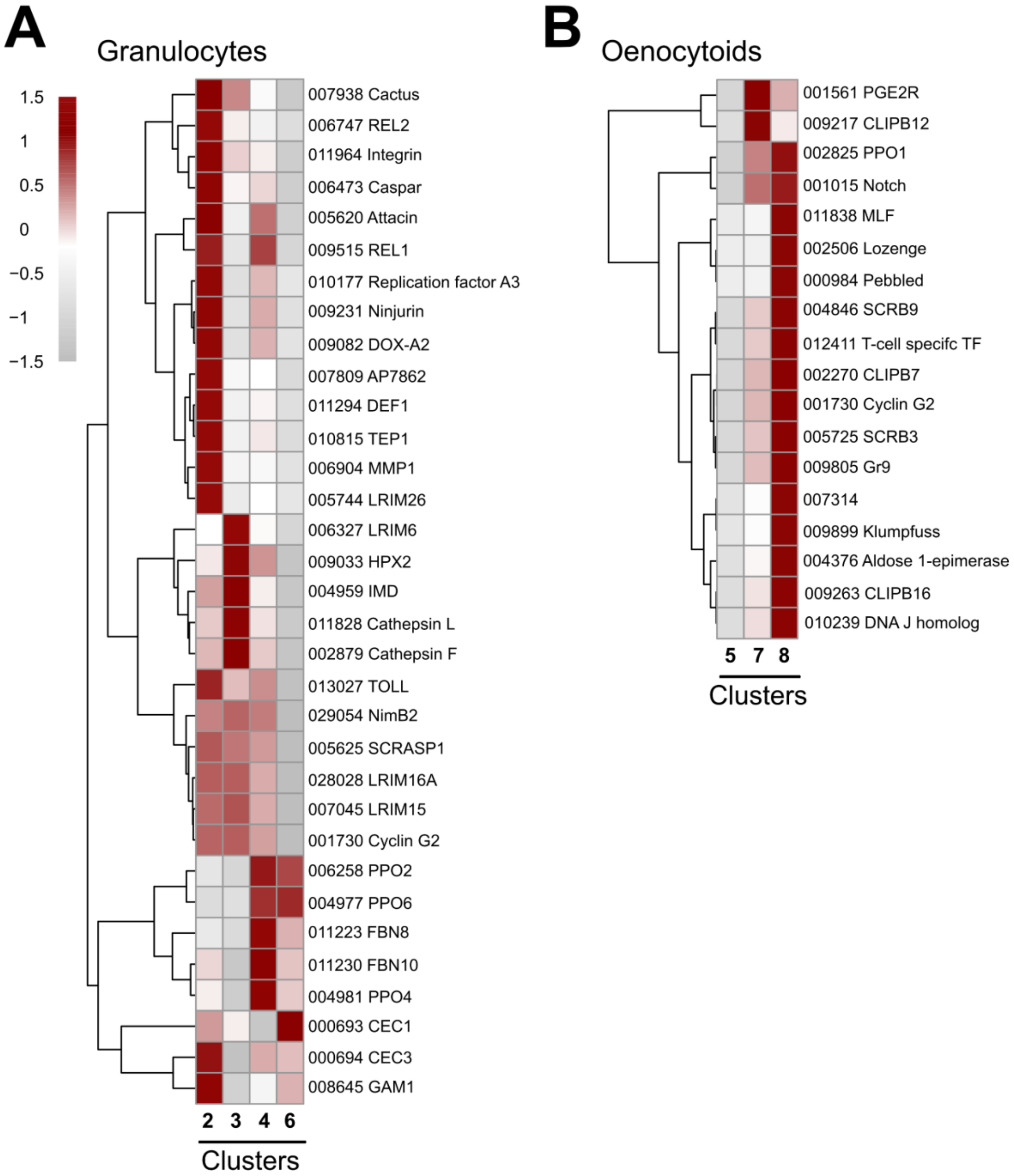


**Figure S13. Expression of granulocyte and oenocytoid markers to distinguish immune cell sub-types.** Expression of marker genes to distinguish granulocyte **(A)** or oenocytoid **(B)** immune cell sub-types. Cluster 5 is included as an outgroup to help distinguish gene expression differences between Clusters 7 and 8 in **(B)**.

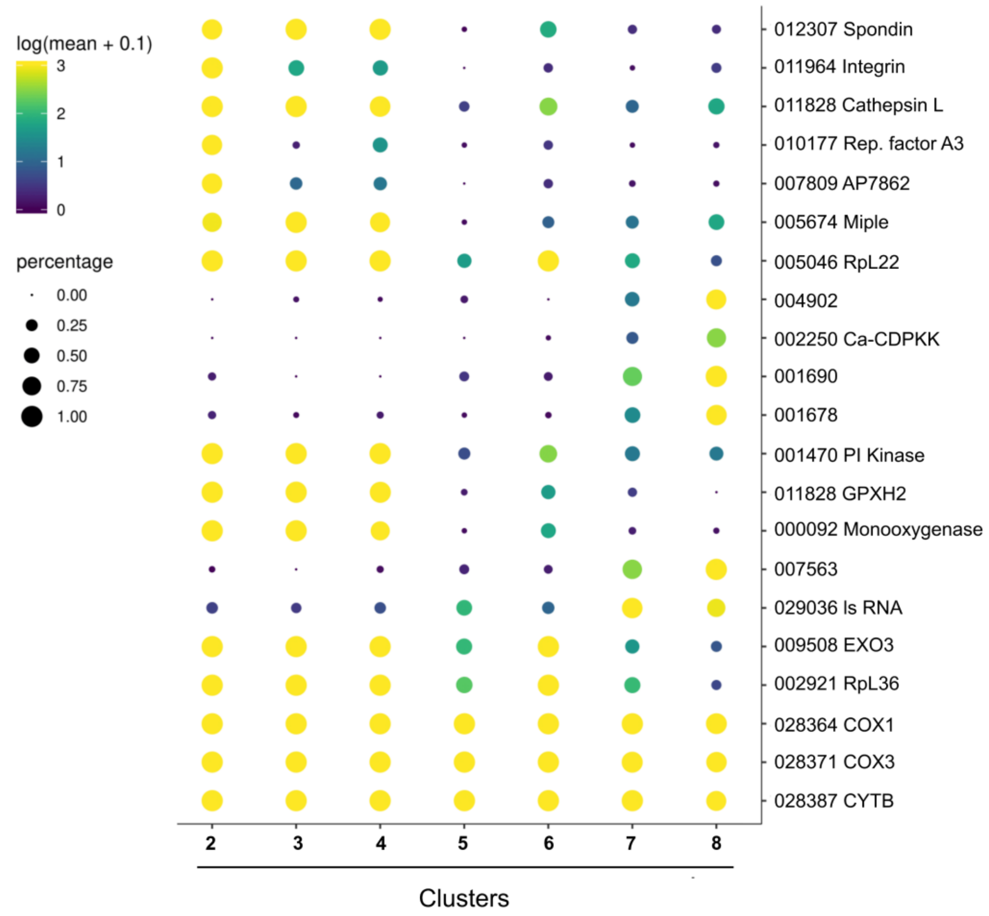


**Figure S14. Marker gene expression across immune cell clusters used to define Monocle3 lineage analysis.** Dot plot displaying the percentage of immune cells and gene expression of candidate marker genes across immune cell clusters. Candidate marker genes are shown on right with AGAP accession number and annotated gene where applicable.

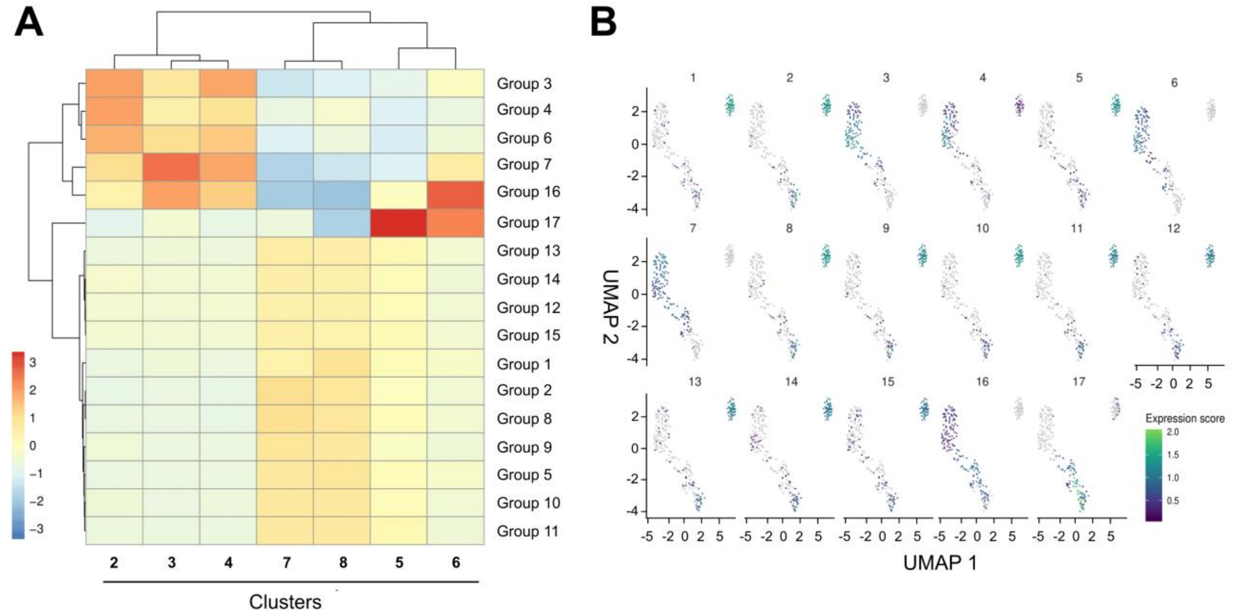


**Figure S15. Enriched transcripts associated with the lineage progressions of mosquito immune cells.** Using Monocle3 analysis, immune cell transcripts were clustered into 17 groups based on their differential expression across immune cell clusters **(A).** Groups are visualized by UMAP displaying cell lineage progressions of related transcripts **(B).**

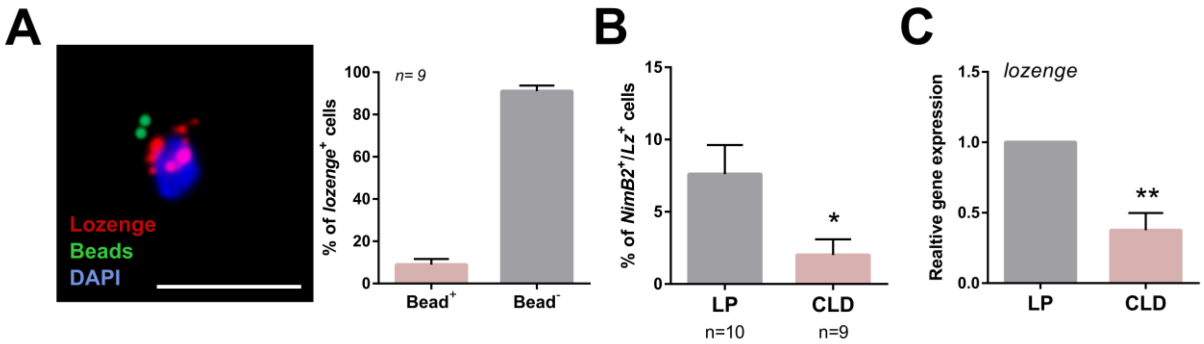


**Figure S16. Phagocytic properties of a subset of *lozenge (lz)*$^+$ cells.** When phagocytosis assays using the injection of fluorescent beads were paired with RNA-FISH experiments, a subset of *lozenge (lz)*$^+$ cells displayed phagocytic properties (bead +) **(A)**. This was confirmed by a reduction in the percentage of *NimB2*$^+$/*Lz*$^+$ cells following treatment with clodronate liposomes (CLD) when compared to liposome only (LP) controls **(B)**, and further validated by qRT-PCR analysis of *lozenge* expression from perfused hemocytes **(C)**. Two independent biological experiments were performed for RNA-FISH experiments in **(A)** and **(B)**. Data were analyzed by Mann–Whitney analysis using GraphPad Prism 6.0. Bars represent mean ± SE. *lozenge* expression was analyzed using an unpaired t test to determine relative gene expression between LP and CLD treatments. Bars represent mean ± SE of three independent replications. Asterisks denote significance (*$P$ < 0.05, ***$P$ < 0.001). Scale bar, 10 µm.

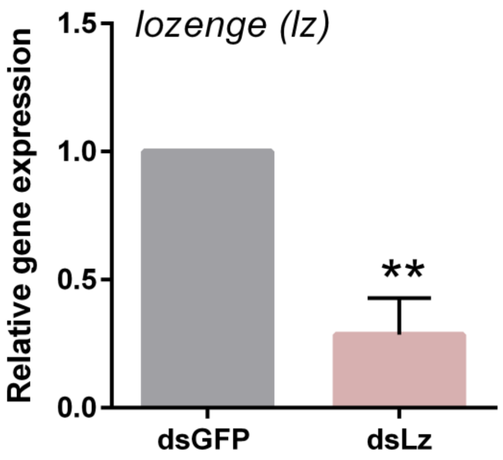


**Figure S17. Validation of *lozenge* knockdown following RNAi.** To disrupt *lozenge* expression, dsRNA corresponding to GFP (control) or lozenge were injected into adult female mosquitoes and evaluated for knockdown 3 days post-dsRNA injection by qRT-PCR. Data from three experiments were analyzed using an unpaired t test in GraphPad Prism 6.0 to determine significance (**, $P<0.01$).
