## Supplementary material for "Single-cell analysis of mosquito hemocytes identifies signatures of immune cell subtypes and cell differentiation": Table S6-Primers for qRT-PCR and dsRNA-mediated gene silencing. (from another version of this preprint: version 1, 2020-07-11)

**Table S6. Primers for qRT-PCR analysis**

**Primer	Gene ID	Sequence (5’- 3’)**

NimB2-F	AGAP029054	CAATCTGCTCAAATGGCTGCTTCCACG

NimB2-R	GCTGCAAACATTCGGTCCAGTGCATTC

LRIM15-F	AGAP007045	CGATCCTGATCCTGAACGTGGGCTTC

LRIM15-R	GCAAGCAAGCCACTCACAAATCCTCG

LRIM16A-F	AGAP028028	ATCAGAGTGCAGCACAAGTTGAAGGT

LRIM16A-R	TCTCTGTTAGCATAGCGCCTTCGTTC

Lz-F	AGAP002506	GCACCGTCAATCAGAACCAA

LZ-R	TGCCACTGATCGAATGCTTG

PPO1-F	AGAP002825	GACTCTACCCGGATCGGAAG

PPO1-R	ACTACCGTGATCGACTGGAC

PPO2-F	AGAP006258	TTGCGATGGTGACCGATTTC

PPO2-R	CGACGGTCCGGATACTTCTT

PPO3-F	AGAP004975	CTATTCGCCATGATCTCCAACTACG

PPO3-R	ATGACAGTGTTGGTGAAACGGATCT

PPO4-F	AGAP004981	GCTACATACACGATCCGGACAACTC

PPO4-R	CCACATCGTTAAATGCTAGCTCCTG

PPO5-F	AGAP012616	GTTCTCCTGTCGCTATCCGA

PPO5-R	CATTCGTCGCTTGAGCGTAT

PPO6-F	AGAP004977	GCAGCGGTCACAGATTGATT

PPO6-R	GCTCCGGTAGTGTTGTTCAC

PPO8-F	AGAP004976	CCTTTGGTAACGTGGAGCAG

PPO8-R	CTTCAAACCGCGAGACCATT

PPO9-F	AGAP004978	TGTATCCATCTCGGACGCAA

PPO9-R	AAGGTTGCCAACACGTTACC

rpS7-F	AGAP010592	ACCCCATCGAACACAAAGTTGACACT

rpS7-R	CTCCGATCTTTCACATTCCAGTAGCAC

SCRB3-F	AGAP005725	CATCGGGACAGCTACATCCT

SCRB3-R	TTATTGCTGCTACCGTTGCC

SCRB9-F	AGAP004846	CGATATTCGGCGATGCAACT

SCRB9-R	CACGCATGACACGATTCAGT
