## Supplementary material for "Single-cell analysis of mosquito hemocytes identifies signatures of immune cell subtypes and cell differentiation": Table S7-Primers for RNAi. (from another version of this preprint: version 1, 2020-07-11)

**Primer	Gene ID	Sequence (5’- 3’)**

T7-GFP F	TAATACGACTCACTATAGGGAGAATGGTGAGCAAGGGCGAGGAGCTGT

T7-GFP R	TAATACGACTCACTATAGGGAGATTACTTGTACAGCTCGTCCATGCC

T7-Lz F	AGAP002506	TAATACGACTCACTATAGGGCTGCAACCGTCCCAGAACAACGGC

T7-Lz R	TAATACGACTCACTATAGGGACAAACCGGAGATCGTTGAATTTGG
